# CardIAP: Calcium images analyzer web application

**DOI:** 10.1101/2021.09.08.452919

**Authors:** Ana Julia Velez Rueda, Agustín García Smith, Luis Alberto Gonano, Maria Silvina Fornasari, Gustavo Parisi, Leandro Matías Sommese

## Abstract

**Motivation:** Ionic calcium (Ca2+) plays the role of the second messenger in eukaryotic cells associated with cellular functions of regulation of the cell cycle, such as transport, motility, gene expression, and metabolism (Permyakov and Kretsinger, 2009). The use of fluorometric techniques in isolated cells, loaded with Ca2+ sensitive fluorescent probes allows the quantitative measurement of dynamic events that occur in living, functioning cells. The Cardiomyocytes Images Analyzer Application (CardIAP) covers the need for tools to analyze and retrieve information from confocal microscopy images, in a systematic, accurate, and fast way.

**Results:** Here we present the CardIAP web app, an automated method for the identification of spatio-temporal patterns in a calcium fluorescence imaging sequence. Through this tool, users can analyze single or multiple Ca2+ transients from confocal line-scan images and obtain quantitative information on the dynamic response of the stimulated myocyte.

Our web application also allows the user the extraction of data on calcium dynamics in downloadable tables and plots, simplifying the calculation of the alternation and discordance indices and their classification. CardIAP could assist in studying the underlying mechanisms of anomalous calcium release phenomena.

**Availability and implementation:** CardIAP is an open-source app, entirely developed in Python, which can be freely accessed and used at http://cardiap.herokuapp.com/.

## Introduction

Cardiovascular diseases (CVD) are the leading cause of global death (GBD, 2018), and in particular, arrhythmias represent a major portion of these deaths (~15–20 %) (Srinivasan and Schilling, 2018). Since abnormal Ca2+ handling is linked to arrhythmias, understanding calcium (Ca2+) management is one of the cardiovascular research’s main goals (Bers, 2014; Landstrom et al., 2017). The Ca2+ release in cardiomyocytes occurs as a consequence of the coordinated opening of multiple Ca2+ release channels (Guatimosim et al. 2002) that can exhibit independent properties and lead to different types of arrhythmogenic events (Hammer et al. 2014). In this sense, image-based techniques could provide valuable spatial and temporal information on cardiomyocytes’ intracellular Ca2+ management (Cannell et al., 1987).

Among the methodologies that offer information about the kinetics of Ca2+ movements more closely to what happens in the cell, confocal microscopy is the most used (Price et al., 2014). However, there are only a few tools available for its analysis (Giovannucci et al., 2019; Schneider et al., 2012), which are usually for general images analysis or studying very specific phenomena. These tools also do not provide the Spatio-temporal analysis of the images, hence this kind of analysis must be carried out through the use of complementary tools (Schneider et al., 2012).

Here we present CardIAP, an open-source web app, that allows the analysis of a series of Ca2+ handling phenomena from a single confocal microscopy image or many. The web interface allows the user to easily manipulate the images of interest to obtain representative amplitude and kinetic data. We aim to provide a user-friendly tool that allows the large-scale analysis of confocal microscopy images.

## Materials and methods

### Calcium Peaks Identification and data processing

CardIAP uses *cardilib*^*1*^ for the image analysis. This python library created by the authors of the present work for performing biomedical images analysis is available for installation and usage in all operative systems through pip.

For analyzing each loaded image, *CardIAP* converts them into a three-dimensional matrix, which corresponds to the size of the image in pixels and the intensity of each pixel. From this matrix, the image is cut in the range selected (ROI) by the user with the cropper displayer and CardIAP analyzes the calcium transients by locating the position of the local maximum intensity.

The algorithm finds the peaks by using the first-order difference (Negri and Vestri, 2017). These potential peaks are filtered using the minimum distance between them, expressed in pixels, and an intensity threshold based on the average of the dataset. The user has the option of retrieving the intensity average across the entire selection over the time series or portioning it and applying the analysis to each of the slices.

CardIAP also provides the user the relative amplitude of each calcium transient, which is calculated from the difference between the maximum and the minimum, divided by the minimum value. To calculate each peak’s rate of decay (tau value), time-series data is cropped from the maximum to the next minimum and applied the least-squares solution to the linear matrix.

### CardIAP implementation

CardIAP is distributed as a web application implemented with the Voilá^2^ framework, which allows the generation of standalone web applications and dashboards from Jupyter notebooks. CardIAP web app takes the images provided by the user as input and allows the images to crop interactively^*3*^ before their analysis. Users also are able to save outputs as CSV tables with the resultant data from both, the complete cell and slices analysis.

CardIAP is available to users through Binder, and it is also hosted^4^ in Heroku’s cloud server. The app’s tutorial and documentation are available at https://cardiap.github.io/, a GitHub page from the original code repository.

### CardIAP access and user interface

Users can upload one or several images from their local disk through the upload button in the CardIAP user interface. After uploading the images, users can optionally crop each image by using the image display, and width and height selector. Once the crop sizes are saved, CardIAP prints them on screen and shows the analysis settings options: slice width, the distance between peaks, and the calibration.

Briefly, the slice width value, which takes values higher than zero (pixels), allows the users to cut the region of interest (ROI) into pieces of the given size in pixels, for calcium locals transients analysis. The distance between peaks value helps CardIAP to enhance the maximum intensity positions threshold. Finally, with the calibration value, users can transform the position of pixels into time data.

The results for each analyzed image will be provided in a different tab. Also, users can check out the information corresponding to all the slices and the complete image analysis separately. CardIAP provides the amplitudes and intensities plots and the downloadable tables with all the analysis results.

### Statistics

A total of 50 images were used for the analysis. Continuous variables were expressed as mean ± SEM and were evaluated with either simple sample Student t-test. To compare the time to peak of both methods, the time data that came out of analyzing the images were normalized with the amplitude. Whenever the distribution was skewed, the Mann-Whitney test was used to compare these data. Pearson’s correlation coefficients were used to measure the relationship between two variables. Levene’s test was used to test the homogeneity of variance of samples. A p-value <0.05 was considered significant.

## Results

For a better understanding of the performance of our software in comparison with existing ones, we made a comparison between amplitude and time-to-peak (TTP) values calculated with CardIAP and the manually extracted data from the same images with an image analyzer program (ImageJ). Figure 2 shows the correlations between both methods. Panel A shows the linear regression that correlates the amplitude values obtained with CardIAP and manually, which presents a Pearson coefficient of 0.98 (p-value <0.0001). Panel B shows the correlation of the TTP normalized values, which presents a Pearson coefficient of 0.89 (p-value <0.0001). From these results, we can infer that both methods yield-related data and the proportion in which one variable estimates the variability of the other.

**Figure 1:**
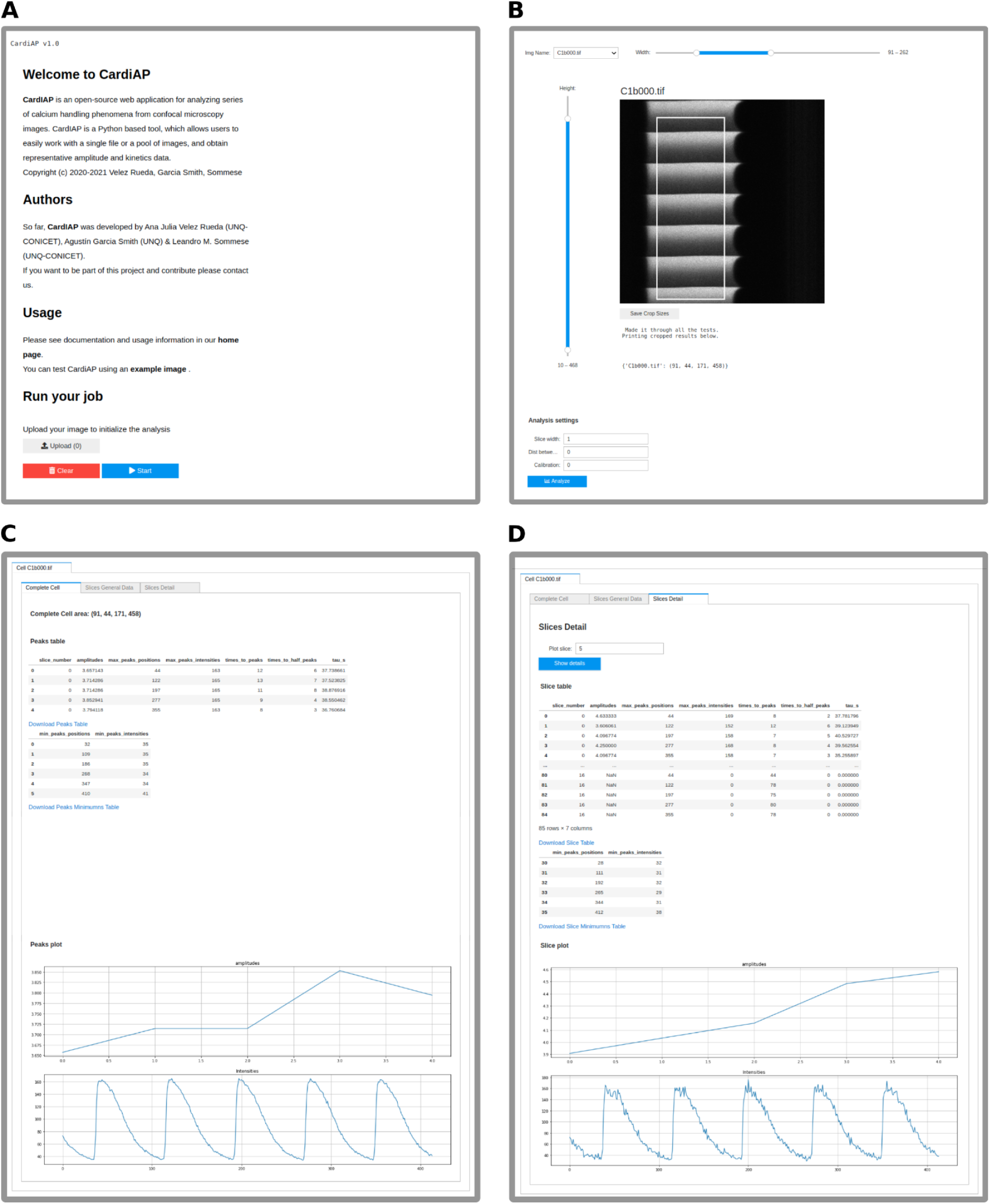
**A**. CardIAP home page. **B**. Once users start the upload of the image panel B displays to allow cropping the image and setting analysis parameters. **C**. This panel shows the first rows of the results of the complete cell analysis. Below each table, there is a link to download them. A graph of average intensity and peak amplitude are displayed across the image to help the user visualize intensity data. **D**. Clicking on the superior label users can visualize the results from different images and the slices of each image.

**Figure 2:**
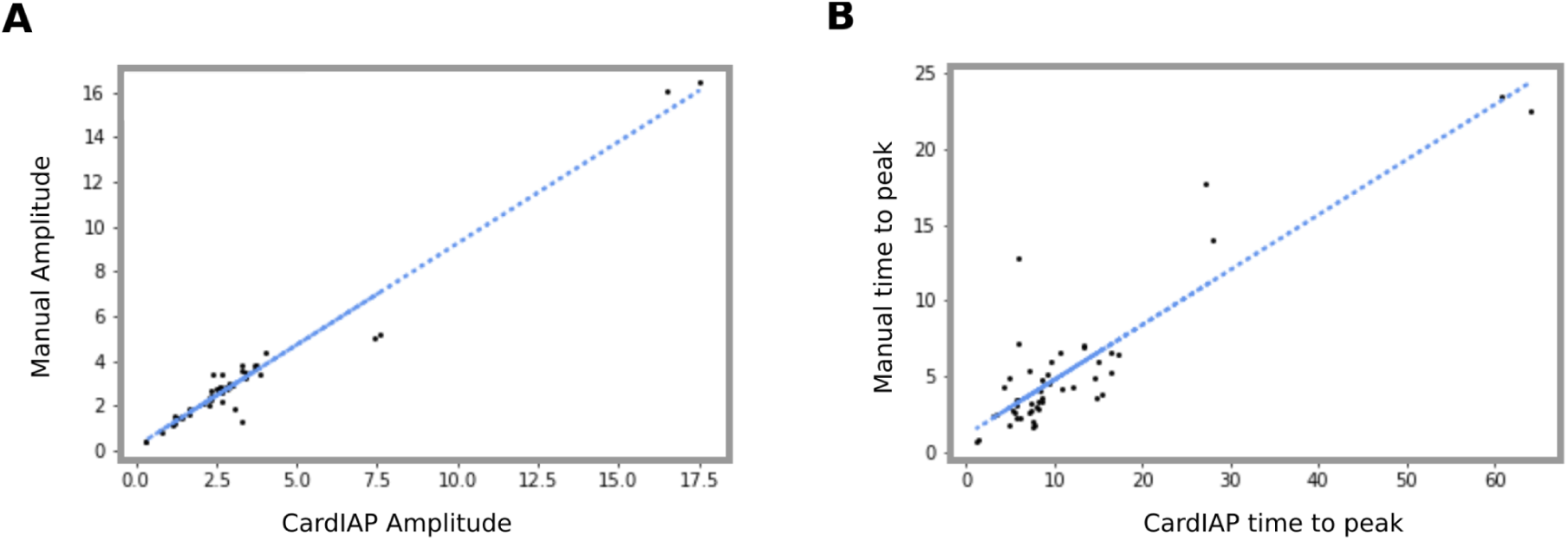
**A**. Correlation plot of the amplitude of calcium transient manually measured and using CardIAP. **B**. Correlation plot of time to peak of calcium transient manually measured and using CardIAP.

To discard whether the measurements related to pixel intensity and time performed with CardIAP show a bias or not, we obtained the paired difference between the values calculated by both methods. For the amplitude measurement, the mean value is 0.12 ± 0.66, and for the TTP 10.36 ± 3.47, which shows that there is no significant bias in amplitude measurement (p-value>0.05), but there is a small bias in the measure of TTP (p-value<0.0001). This observation could be explained by the dispersion and distribution of the measures, as can be observed by comparing the standard deviation in both methods. The manual measurement takes values with greater variance concerning the average values (variance: 7.96, mean: 10.10), contrary to what happens with CardIAP measurement (variance: 4.17, mean: 20.79)

## Conclusion

Here we presented CardIAP, a free web application, that aims to ease the analysis of multiple line scan images taken by calcium transient microscopy techniques. Our web application transforms cell images in time series plots to accurately detect transient peaks, using our analysis library cardilib. CardIAP measures the intensity and kinetics parameters of the peaks quickly and transparently so that the user can have control of the processes that occur automatically.

CardIAP offers the user the possibility of analyzing a set of images in parallel but selecting the area of interest individually. This, in contrast with manual image processing, allows the study to be replicable and systematic, reducing bias in reading the plots. Therefore, CardIAP prevents the user from using multiple tools, reducing the analysis time and the resources to be used. Furthermore, as explained above, CardIAP gives users the possibility of dividing each image into portions and replicating the obtaining of characteristic parameters of the calcium transients for each of them.

The easy way of retrieving calcium dynamics data through CardIAp facilitates its application not only to study calcium transients but also other calcium releases phenomena (Edwards and Blatter, 2014; Hohendanner et al., 2016; Skardal et al., 2014), such as calcium waves and sparks, or to study anomalous calcium release phenomena such as alternation, discordance, and dyssynchrony.

## Footnotes

1 Cardilib: https://pypi.org/project/cardilib/

2 Voilá: https://voila.readthedocs.io/en/stable/index.html

3 *Interactivecrop*: https://pypi.org/project/interactivecrop/

4 CardiAP: http://cardiap.herokuapp.com/

## References

Bers, D.M. (2014). Cardiac sarcoplasmic reticulum calcium leak: basis and roles in cardiac dysfunction. Annu. Rev. Physiol. 76, 107–127.

Cannell, M.B., Berlin, J.R., and Lederer, W.J. (1987). Intracellular calcium in cardiac myocytes: calcium transients measured using fluorescence imaging. Soc. Gen. Physiol. Ser. 42, 201–214.

Edwards, J.N., and Blatter, L.A. (2014). Cardiac alternans and intracellular calcium cycling. Clin. Exp. Pharmacol. Physiol. 41, 524–532.

GBD (2018). Global, regional, and national age-sex-specific mortality for 282 causes of death in 195 countries and territories, 1980-2017: a systematic analysis for the Global Burden of Disease Study 2017. Lancet 392, 1736–1788.

Giovannucci, A., Friedrich, J., Gunn, P., Kalfon, J., Brown, B.L., Koay, S.A., Taxidis, J., Najafi, F., Gauthier, J.L., Zhou, P., et al. (2019). CaImAn an open source tool for scalable calcium imaging data analysis. Elife 8.

Hohendanner, F., DeSantiago, J., Heinzel, F.R., and Blatter, L.A. (2016). Dyssynchronous calcium removal in heart failure-induced atrial remodeling. Am. J. Physiol. Heart Circ. Physiol. 311, H1352–H1359.

Landstrom, A.P., Dobrev, D., and Wehrens, X.H.T. (2017). Calcium signaling and cardiac arrhythmias. Circ. Res. 120, 1969–1993.

Negri, L.H., and Vestri, C. (2017). Lucashn/Peakutils: V1.1.0. Zenodo.

Permyakov, E.A., and Kretsinger, R.H. (2009). Cell signaling, beyond cytosolic calcium in eukaryotes. J. Inorg. Biochem. 103, 77–86.

Price, R.L., Haley, S.T., Bullard, T., Davis, J., Borg, T.K., and Terracio, L. (2014). Confocal microscopy of cardiac myocytes. Methods Mol. Biol. 1075, 185–199.

Schneider, C.A., Rasband, W.S., and Eliceiri, K.W. (2012). NIH Image to ImageJ: 25 years of image analysis. Nat. Methods 9, 671–675.

Skardal, P.S., Karma, A., and Restrepo, J.G. (2014). Spatiotemporal dynamics of calcium-driven cardiac alternans. Phys. Rev. E Stat. Nonlin. Soft Matter Phys. 89, 052707.

Srinivasan, N.T., and Schilling, R.J. (2018). Sudden cardiac death and arrhythmias. Arrhythm. Electrophysiol. Rev. 7, 111–117.

